## Supplementary Information for "Deep Reinforcement Learning for Optimal Experimental Design in Biology"

### 1 Fitted Q-learning can design optimal experiments on a simple Monod growth system

To test the potential of using reinforcement learning for OED, we first applied the Fitted Q-learning algorithm to a simple non-linear system with Monod dynamics to investigate whether the agent is capable of optimising the D-optimality score with the given information. We compare the performance of the FQ agent with a one step ahead optimiser (OSAO). In this system we have one state variable, $x$, and no measurement noise so that output $\mathbf{Y} = x$. There is one control variable $u$ that is controlled by the OSAO or the RL controller. The dynamics of the system are given by a simple Monod relationship between $\frac{dx}{dt}$ and $u$:

$$\frac{dx}{dt} = \frac{p_1 u}{p_2 + u} x,$$

where $p_1$ and $p_2$ are the parameters to be estimated.

In the following $p_1 = p_2 = 1$. Both the RL and OSAO implementations start from initial condition $x_0 = 1$, $u_0 = 0.5$ and are able to choose $u$ between $0 \leq u \leq 0.1$. The FQ agent works in a discrete action space and therefore has ten equally spaced discrete actions to choose from (distributed uniformly between 0 and 0.1). Figure S1A shows the experimental input profiles chosen by the FQ agent and OSAO, which are similar. These consist of inputs at the highest level available and a value about half way between the maximum and minimum values. The FQ agent selects inputs values that straddle the optimum found by the OSAO, likely due to the discrete nature of its action space. Figure S1B shows the system trajectories of both controllers. As expected these are very similar. Figure S1C shows the performance of the FQ agent as it was trained for 500 episodes, compared to the performance of the trajectory found by the OSAO. Their performance is similar; by the end of training, the FQ agent is performing slightly better than the OSAO.

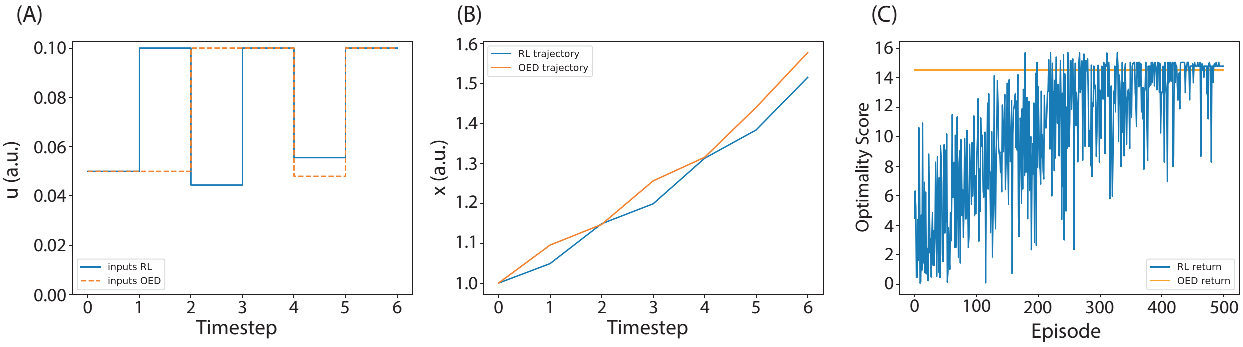


Figure 1: Reinforcement learning for optimal experimental design on a simple non-linear system. (A) The input profiles of the two agents. (B) The corresponding state trajectories. (C) The training performance of the RL agent compared to the performance achieved by the OSAO.

### 2 Full value fitting results

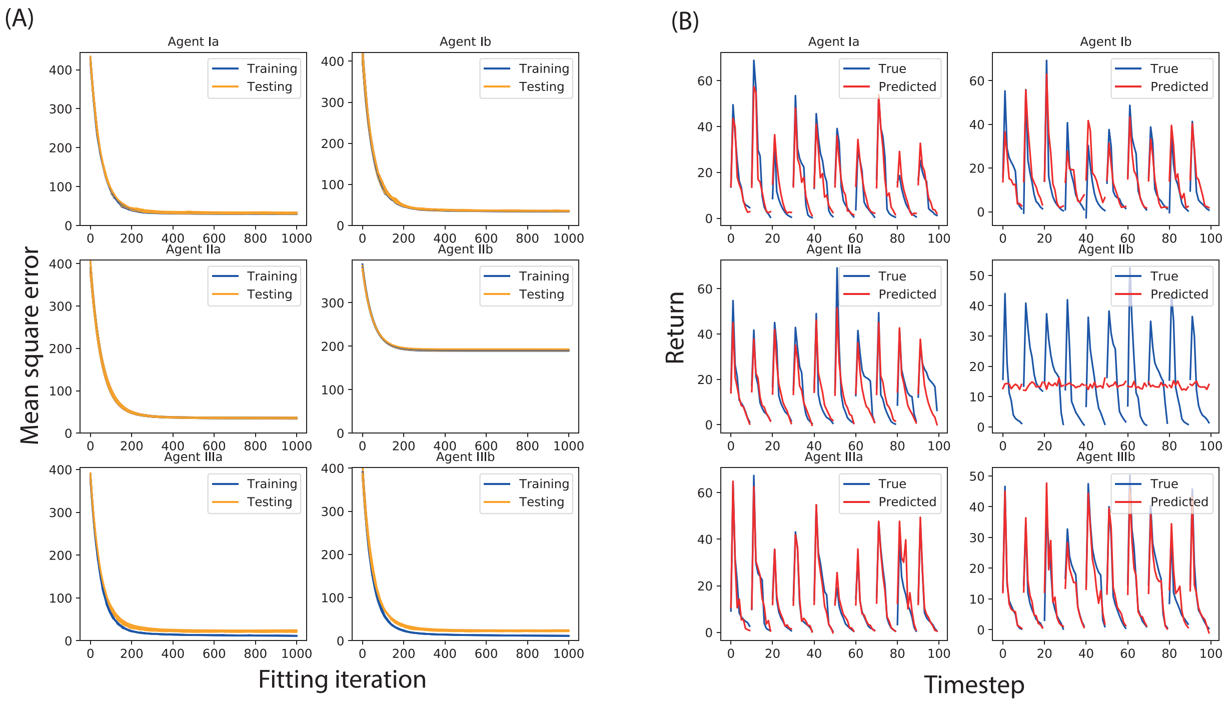


Figure 2: Full value fitting results (A) Training and testing performance of all agents over 1000 Fitted Q-iterations. (B) The true vs predicted return of 10 experiments for all agents after training.

#### 3 Recurrent FQ

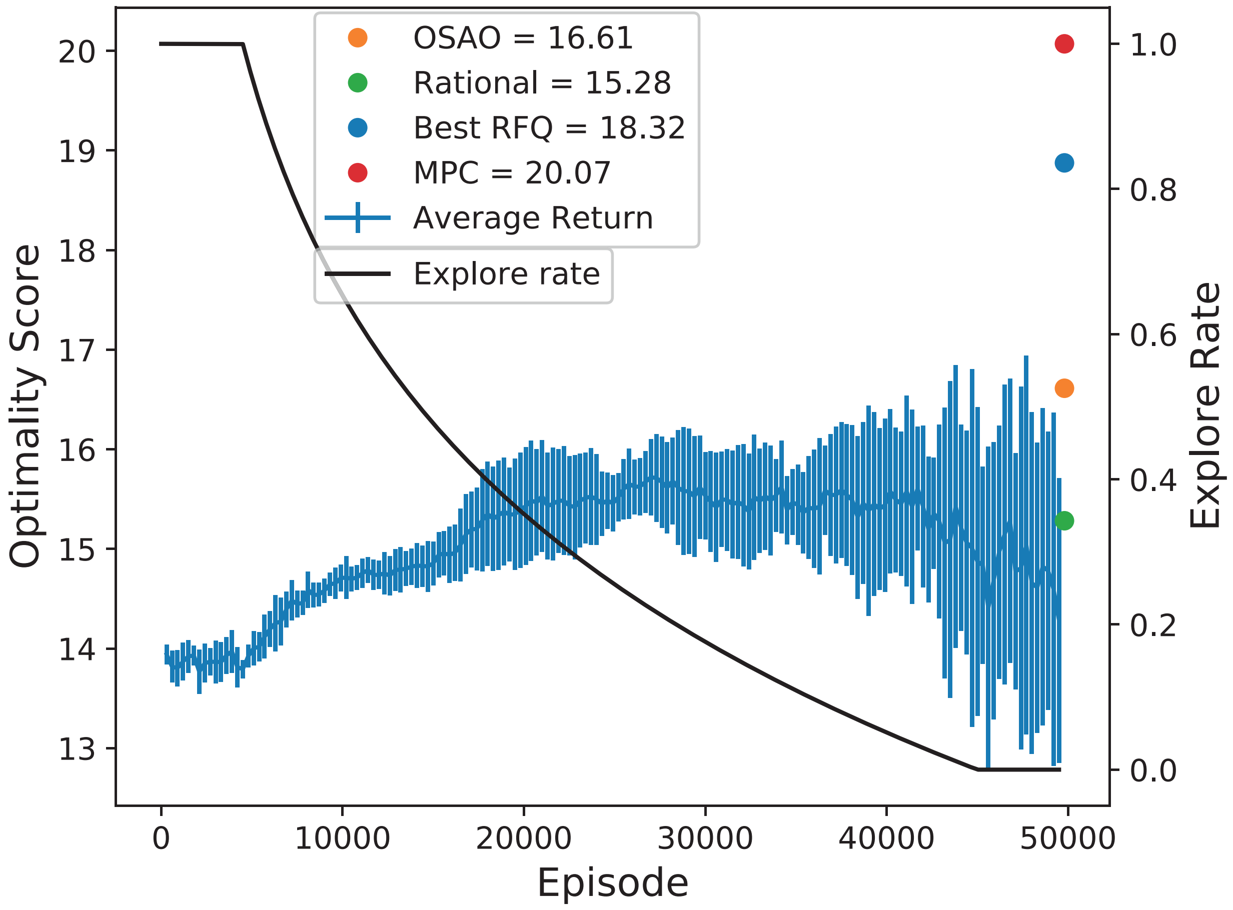


Figure 3: Recurrent FQ for optimal experimental design to infer the values of model parameters for an auxotrophic bacterial strain growing in a chemostat. Average training progress of nine recurrent FQ-agents over 50000 episodes. The return of the FQ-agents is averaged across the 9 repeats. The mean is shown, along with error bars indicating one standard deviation.

### 4 RT3D performance for different parameter samples

| Parameter value $[\mu_{max}, K_m, K_{m0}]$ | MPC | RL |
|---|---|---|
| $[0.552564, 0.000400962, 0.0000775143](S1)$ | 18.85 | 17.63 |
| $[0.708972, 0.000500437, 0.0000490196](S2)$ | 20.05 | 19.03 |
| $[0.500478, 0.00041873, 0.000029529](S3)$ | 18.02 | 16.81 |
| $[1.45073, 0.000810734, 0, 0000961402](S4)$ | 21.52 | 20.41 |
| $[0.5, 0.0001, 0.00001]$ (Lower bound) | 18.01 | 16.78 |
| $[1.25, 0.00055, 0.000055]$ (Centre) | 20.79 | 20.31 |
| $[1, 0.00048776, 0.00006845928]$ (Nominal params) | 20.07 | 20.11 |
| $[2, 0.001, 0.0001]$ (Upper bound) | 21.86 | 20.15 |

Table 1: Comparison of RL OED controller trained over a parameter distribution compared with an MPC with perfect system knowledge. The optimality score of the experiments produced by each controller is shown for different samples within the distribution.
